# Honeybee Queen mandibular pheromone induces starvation in *Drosophila melanogaster*, implying a role for nutrition signalling in the evolution of eusociality

**DOI:** 10.1101/2021.04.08.439099

**Authors:** Mackenzie R. Lovegrove, Elizabeth J. Duncan, Peter K. Dearden

**Affiliations:** Genomics Aotearoa and Biochemistry Department, University of Otago, P.O. Box 56, Dunedin, Aotearoa, New Zealand; School of Biology, Faculty of Biological Sciences, University of Leeds, Leeds LS2 9JT, UK

## Abstract

Eusocial insect societies are defined by the reproductive division of labour, a social structure that is generally enforced by the reproductive dominant or ‘queen’. Reproductive dominance is maintained through behavioural dominance in some species as well as production of queen pheromones in others, or a mixture of both. Queen mandibular pheromone (QMP) is produced by honeybee (*Apis mellifera*) queens and has been characterised chemically. How QMP acts to repress worker reproduction, and how it has evolved this activity, remains less well understood. Surprisingly, QMP is capable of repressing reproduction in non-target arthropods which have not co-evolved with QMP, are never exposed to QMP in nature, and are up to 530 million years diverged from the honeybee.

Here we show that, in *Drosophila melanogaster*, QMP treatment mimics nutrient limiting conditions, leading to disrupted reproduction. Exposure to QMP induces an increase in food consumption, consistent with that observed in *D. melanogaster* in response to starvation conditions. This response induces the activation of two checkpoints within the ovary that inhibit oogenesis. The first is the 2a/b ovarian checkpoint in the germarium, which reduces the flow of presumptive oocytes. A stage 9 ovarian checkpoint is also activated, causing degradation of oocytes. The magnitude of activation of both checkpoints is indistinguishable between QMP treated and starved individuals.

As QMP seems to trigger a starvation response in an insect highly diverged from honeybees, we propose that QMP originally evolved by co-opting nutrition signalling pathways to regulate reproduction, a key step in the evolution of eusociality.

## Introduction

In eusocial animal societies individuals in a group are divided between non-reproductive and reproductive roles. Reproductively dominant individuals, often called queens, are the primary reproductive individuals whereas workers carry out all the other tasks in the group (Michener, 1974; Oster and Wilson, 1978). Eusociality has evolved independently at least 16 times within the insects (Crespi, 1992; Grimaldi and Engel, 2005; Inward et al., 2007; Kent and Simpson, 1992; Tanaka and Itô, 1994), and up to 11 times in the Hymenoptera-containing the bees, ants and wasps (Brady et al., 2006; Cameron and Mardulyn, 2001; Crozier, 2008; Danforth et al., 2013; Hines et al., 2007; Hughes et al., 2008; Moreau et al., 2006; Peters et al., 2017).

Queens often regulate their own colony by producing chemical cues (queen pheromones) which actively repress the reproduction of her female subordinate workers. Perhaps the most well-studied queen pheromone is queen mandibular pheromone (QMP) produced by the queen honeybee, *Apis mellifera* (Keeling et al., 2003; Pankiw et al., 1996; Pham-Delègue et al., 1993; Princen et al., 2019). QMP is chemically distinct from other social hymenopteran queen pheromones, including those seen in *Bombus terrestris* (Van Oystaeyen et al., 2014) and it has presumably undergone significant evolution in the time period since these species diverged 55 million years ago (Peters et al., 2017).

QMP not only has the ability to repress reproduction in worker honeybees, but in non-target arthropods as well. It has been shown to repress reproduction in *Drosophila melanogaster* (Camiletti et al., 2016; Lovegrove et al., 2019; Sannasi, 1969), a house fly (*Musca domestica*) (Nayar, 1963), bumble bee (*Bombus terrestris*) (Princen et al., 2020), an ant (*Formica fusca*) (Carlisle and Butler, 1956), termite (*Kalotermes flavicollis*) (Hrdy et al., 1960) and a prawn (*Leander serratus*) (Carlisle and Butler, 1956); species that shared a last common ancestor more than 530 million years ago (dos Reis et al., 2015). This ability to repress reproduction in a broad range of species is a novel feature, unique to QMP, and not a conserved function of hymenopteran queen pheromones (Lovegrove et al., 2019).

The phylogenetically broad effect of QMP, despite its relatively recent evolution is a paradox. Honeybee QMP contains a mixture of chemicals that differ from bumble bee and other hymenopteran queen pheromones, implying significant evolution of this pheromone since bumblebees and honeybees diverged. This relatively recent evolution has led to QMP having an activity that represses reproduction in species which diverged from the honeybee lineage hundreds of millions of years before QMP evolved; an activity not found in other hymenopteran queen pheromones (Lovegrove et al., 2019). One possible explanation for this is that QMP may interact with pathways that are conserved across arthropods. This implies that examination of how QMP works to repress reproduction in tractable model systems may help us understand its activity in honeybees.

Female *Drosophila melanogaster*, while being 340 million years diverged from the honeybee (Misof et al., 2014), are consistently repressed by exposure to QMP (Camiletti et al., 2013; Lovegrove et al., 2019; Sannasi, 1969). *D. melanogaster* therefore provide a tool to uncover the mechanisms by which QMP acts to repress reproduction, thus informing future work in the target species, *Apis mellifera*.

In this study, we have investigated the mechanisms by which QMP represses reproduction in *D. melanogaster*. We have discovered that honeybee queen pheromone acts in this non-target species by inducing a starvation response, leading to alterations in food consumption and the activation of ovarian checkpoints consistent with starvation-induced repression. In doing so, we have demonstrated for the first time a mechanism by which QMP acts in a non-target species which may provide insight into the evolution of QMPs activity.

## Results

### QMP repression in the Drosophila ovary is not acting through loss of ovarioles

The *D. melanogaster* ovary is comprised of an average of 15 ovarioles per ovary-the chains along which oocytes develop and mature (King, 1970; Wu et al., 2008). The anterior-most region is the germarium, containing the germline cells. Within this region, egg chambers develop (Wu et al., 2008), which leave the germarium, and progress along 14 stages of development, resulting in the production of a mature oocyte (King, 1970).

One mechanism by which QMP is reported to reduce reproduction in worker honeybees is by reducing the number of ovarioles (Ronai et al., 2017) thus reducing reproductive potential. To investigate how QMP is reducing the reproductive capacity of female *D. melanogaster*, we first investigated whether the reduction in the number of mature oocytes produced was due to a loss of entire ovarioles. The number of germaria per ovary were counted for individuals which had been exposed to QMP or a solvent control for 12, 24 or 48 h. Also included was a starved control, as the phenotype of starvation is well-established in the *D. melanogaster* ovary (Pritchett et al., 2009). Across time and treatment, the number of germaria per ovary ranged from an average of 12.99-14.76 (Fig 1A). We observed no significant differences in ovariole number between any of the treatments (AOD χ^2^ = 1.2457, df = 2, *p* = 0.5364), including at 48 hours, a timepoint at which we see a 70% reduction in the number of mature eggs (Figure 1B), suggesting that QMP does not impair reproductive capacity in *D. melanogaster* by reducing the number of ovarioles within an ovary.

**Figure 1.**
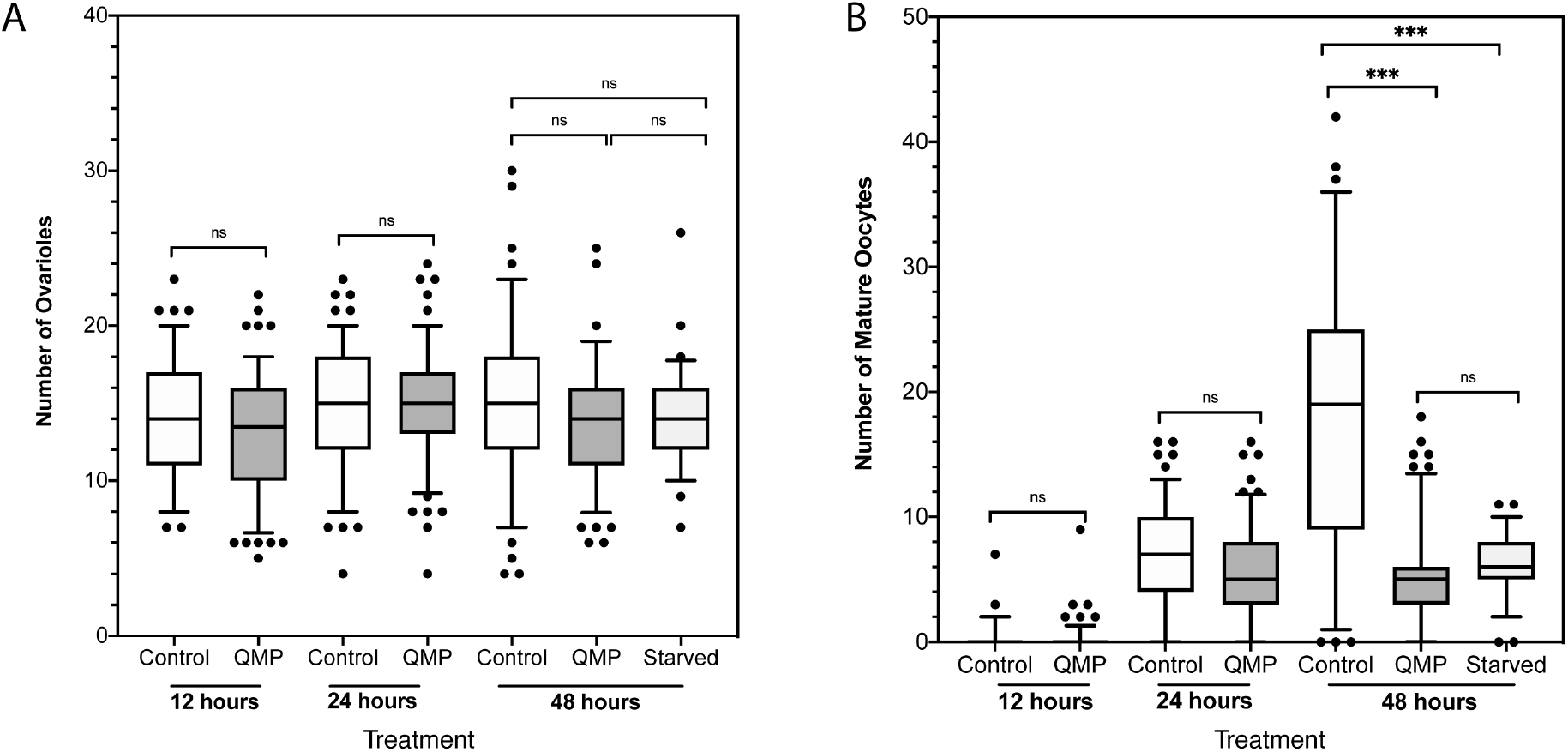
**A)** Box and whisker plot showing the number of ovarioles per ovary in *D. melanogaster* that were exposed to either 26 Qe QMP or an ethanol solvent control for 12, 24 or 48 h. Also included was a 48 h starved positive control. Statistical analysis was carried out using a generalised linear mixed model (GLMM) with a Poisson distribution, there was no statistically significant differences in ovariole number between any of the treatments. **B)** Box and whisker blot showing the average number of mature (stage 14 oocytes) per ovary in *D. melanogaster* exposed to either 26 Qe QMP or an ethanol solvent control for 12, 24, 36 or 48 h. Also included was a 48 h starvation control. Differences in mature oocyte number were determined using a GLMM with a negative binomial distribution followed by Tukey post-hoc analysis, statistically significant differences are by asterisks (***, p < 0.001). In both A and B the box is defined by 25th percentile, median and 75th percentile, whiskers extend to 5% and 95%; and outliers are represented by individual points.

After establishing that QMP does not repress reproduction by reducing the number of ovarioles, we then investigated the end product of reproduction, the number of mature oocytes (stage 14) which were being produced. Virgin females were exposed to 26 Qe QMP or a solvent control for 12 h – 48 h. After exposure, their ovaries were dissected out and the number of mature oocytes counted as a measure of fecundity.

A significant difference in the number of mature oocytes between QMP exposed and control individuals was observed was after 48h (AOD χ^2^ = 17.689, df = 2, *p* = 0.1441 × 10^−3^). After 12 h (control mean = 0.2353, QMP mean = 0.2256, *p =* 1.00) and 24 h (control mean = 3.850, QMP mean = 3.720, *p =* 0.5268) of exposure there was no observable difference in the number of oocytes. The number of mature oocytes produced by the controls, however, increased steadily across the time course (Fig. 1B, 12 h mean = 0.2353, 24 h mean = 7.331 48 h mean = 17.820). Those exposed to QMP, however, produced approximately 5 mature oocytes after 24 hours of exposure (12 h mean = 0.2256, 24 h mean = 5.553) - a number which remained static between 24 h and 48 h (48h mean = 5.015) (Fig 1B). QMP exposure did, however, result in a significant difference in the number of mature oocytes between controls and treated individuals (control mean = 17.820, QMP mean = 5.015, *p <* 0.0001). Starvation also caused a decrease in the number of mature oocytes produced (control mean = 17.820, starvation mean = 6.172, *p <* 0.0001), reducing mature oocytes to a similar level observed with QMP exposure (QMP mean = 5.015, starvation mean = 6.172 *p <* 0.8481).

### QMP activates the 2a/b ovarian checkpoint to reduce fecundity consistent with a starvation response

QMP acts in *D. melanogaster* to reduce the number of mature oocytes produced, not to reduce the number of ovarioles (Fig. 1). We next tested if the loss of mature oocytes was due to the activation of checkpoints earlier in oogenesis. Much is already known about how reproduction is controlled in *D. melanogaster* in response to environmental stimuli (Pritchett et al., 2009). Regulation of reproduction in *D. melanogaster* is associated with the action of ovarian checkpoints, which regulate oogenesis and oocyte maturation.

The first of these is located in the germarium at stage 2a/b (Fig. 2A), where the follicle cells begin to surround a germ line cyst, and polarisation of the oocyte begins (Huynh and St Johnston, 2004). Nutrient deprivation leads to an increase in apoptosis in this region and a decrease in oocyte production (Drummond-Barbosa and Spradling, 2001). The second, is located in mid-oogenesis, at stage 9 characterised by the anterior follicle cells beginning to migrate to the posterior and encapsulate the oocyte. When this checkpoint is activated, the stage 9 oocytes undergo cell death. This is characterised by the fragmentation and condensation of nuclei (McCall, 2004).

**Figure 2.**
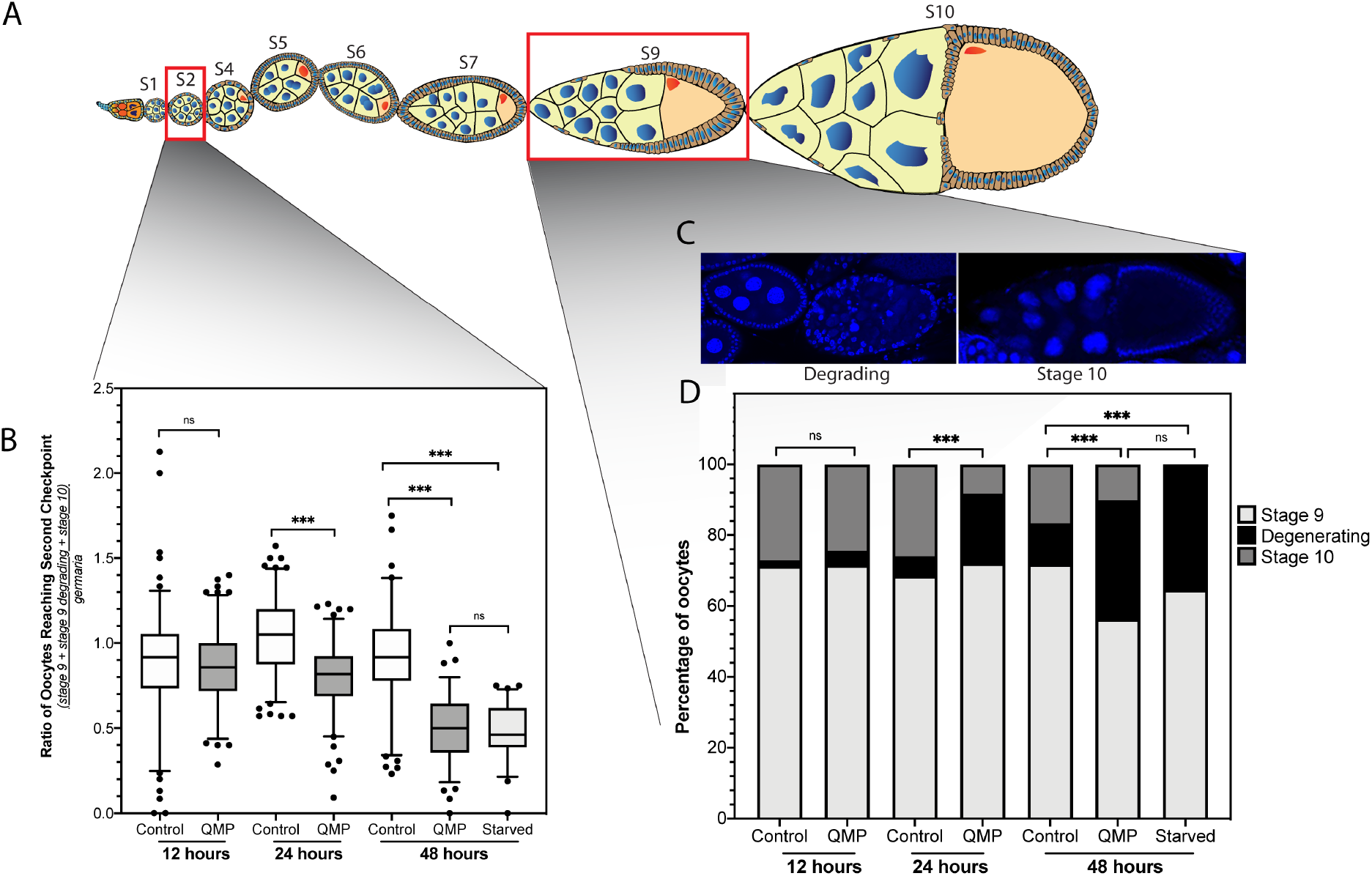
A) Schematic of a *Drosophila* ovariole with stages and positions of the two ovarian checkpoints marked B) Ratio of germaria compared to the number of oocytes which are reaching or passing through the stage 9 ovarian checkpoint. *D. melanogaster* were exposed to either a control or 26 Qe QMP for 12, 24 or 48 hours. Also included was a 48 hr starved control. The box is defined by 25th percentile, median and 75th percentile, whiskers extend to 5% and 95%; the outliers are represented with circles. Differences in oocytes that had passed the Stage 2 checkpoint were determined using a GLMM with a Gaussian distribution followed by Tukey post-hoc analysis, statistically significant differences are by asterisks (***, p < 0.001). C) Representative images of stage 9 degenerating and stage 10 oocytes used to calculate the effect of the stage 9/10 checkpoint. D) Stacked bar chart shows the proportion of oocytes in *D. melanogaster* ovaries at stage 9, stage 9 degenerating and stage 10. Individuals were exposed to 26 Qe QMP or an ethanol solvent control for 12, 24 or 48 hours. Also included is a positive control for 48 h of starvation. As the number of ovarioles, stage 9 oocytes, degenerating oocytes and stage 10 oocytes are likely dependent on each other, data was analysed by performing a principal components analysis (PCA), which indicated that the majority of variation was observed in the first principal component. This component was extracted and a GLMM with a Gaussian error structure and a Tukey post-hoc test was used to determine the effect of treatments at different time points. Statistically significant differences are by asterisks (***, p < 0.001).

We found that QMP was reducing the number of oocytes which progressed past the first checkpoint (stage 2a/b) to the second checkpoint (Fig. 2A, AOD χ^2^ = 33.265, df = 2, *p* = 5.977 × 10^−8^). This reduction was not observed after 12 h of QMP exposure (the germaria produced a ratio of 0.878 oocytes reaching the stage 9 checkpoint in the control and 0.872 oocytes reaching the stage 9 checkpoint in QMP exposed individuals, *p =* 1). After 24 h of QMP exposure, however, there was a significant difference as controls had, on average, 1.032 oocytes reaching stage 9 per germarium, whereas the QMP treated flies had only 0.797, a 23% reduction (*p=* 0.0005). This difference became more pronounced by 48 hours of exposure, where controls produced 0.945 stage 9 oocytes per germaria, and the QMP exposed had 0.505 (*p=* <.0001). Therefore, QMP exposure reduced the number of oocytes passing through the 2a/b checkpoint by 46.56% after 48 h of exposure. There was no significant difference between individuals which had been exposed to QMP for 48 h (QMP exposed mean 0.505) and those starved for 48 h (Starvation mean = 0.477) (*p* = 1.00). This indicates that QMP acts to repress reproduction in *D. melanogaster* by reducing the flow of oocytes through the 2a/b ovarian checkpoint by half, similar to starvation.

### QMP activates the stage 9 ovarian checkpoint to reduce fecundity consistent with starvation

We have shown evidence of a QMP triggered checkpoint (stage 2a/b) acting after the germarium, but prior to the stage 9 ovarian checkpoint (Fig. 2A). Activation of the stage 9 checkpoint would result in a reduction of the number of mature oocytes. Therefore, we investigated whether the stage 9 checkpoint itself was activated in response to QMP exposure (Fig. 2D). This was carried out by counting the number of stage 9 oocytes, stage 9 degenerating and stage 10 oocytes. This data is shown proportionally to remove the effect of the stage 2a/b checkpoint; we have shown the stage 2a/b checkpoint is significantly activated in response to QMP exposure (Fig. 2B), which reduces the number of oocytes that reach the stage 9 checkpoint. To account for the differences in activation of the early checkpoint we compare the proportions of oocytes reaching stage 9, degrading or passing the checkpoint to reach stage 10 to determine if QMP exposure is also affecting the stage 9 checkpoint (Fig. 2D)

Our data indicates a significant effect of time and treatment on the numbers of stage 9, degenerating and stage 10 oocytes (Fig. 2B, AOD χ^2^ = 17.191, df = 2, *p* = 1.849 × 10^−4^). There was no significant difference observed in the proportion of oocytes after 12 h QMP exposure (*p =* 0.7725) but by 24 h of QMP exposure, there was a significant different in the proportions of oocyte stages (*p* = <0.0001). This difference was not observed in the number of healthy oocytes at stage 9 (controls had 68.56% and QMP treated has 72.14%), but instead in the proportion of oocytes degenerating at stage 9. Controls had 5.41% of their oocytes degenerating at stage 9, whereas the QMP treated had 19.62%-an almost 4-fold increase. This degeneration resulted in a decrease in the number of oocytes seen at stage 10, where controls had 26.04% and QMP treated had just 8.24%.

After 48 h of exposure to QMP there was a continued significant difference in the proportions of oocytes seen at each stage (*p* = <0.0001; at this time point there were less oocytes reaching stage 9 (Control mean = 71.83% and QMP treated mean = 56.30%) consistent with activation of the stage 2a/b ovarian checkpoint (Fig. 2A). At 48 h 11.53% of the control oocytes were degenerating, versus 33.54% of the QMP treated. This degeneration resulted in a decrease in the number of oocytes seen at stage 10, where controls had 16.64% and QMP treated had just 10.16%.

Intriguingly there was no significant difference in the proportion of oocytes seen at stage 9, stage 9 degenerating or stage 10 when the 48 h starved and 48 h QMP were compared (*p* = 1.000), implying that the activation of the second ovarian checkpoint is similar between QMP and starvation treatments.

### QMP exposed flies consume more food in a pattern consistent with starvation

We have shown that QMP treatment causes activation of both the stage 2a/b and stage 9 checkpoints in the *D. melanogaster* ovary (Fig. 2), a phenotype similar to that caused by starvation. To test the hypothesis that QMP treatment may be mimicking starvation we carried out a food intake assay. *D. melanogaster* were exposed to QMP, or an ethanol control, for 48 h with no restrictions on food intake before being allowed to consume coloured food solution *ad libitum* for 2 h. Their feeding behaviour was determined by examining their abdomens and scoring them based on the proportion of the abdominal area that was coloured (Jiang et al., 2018) (Fig. 3A).

**Figure 3.**
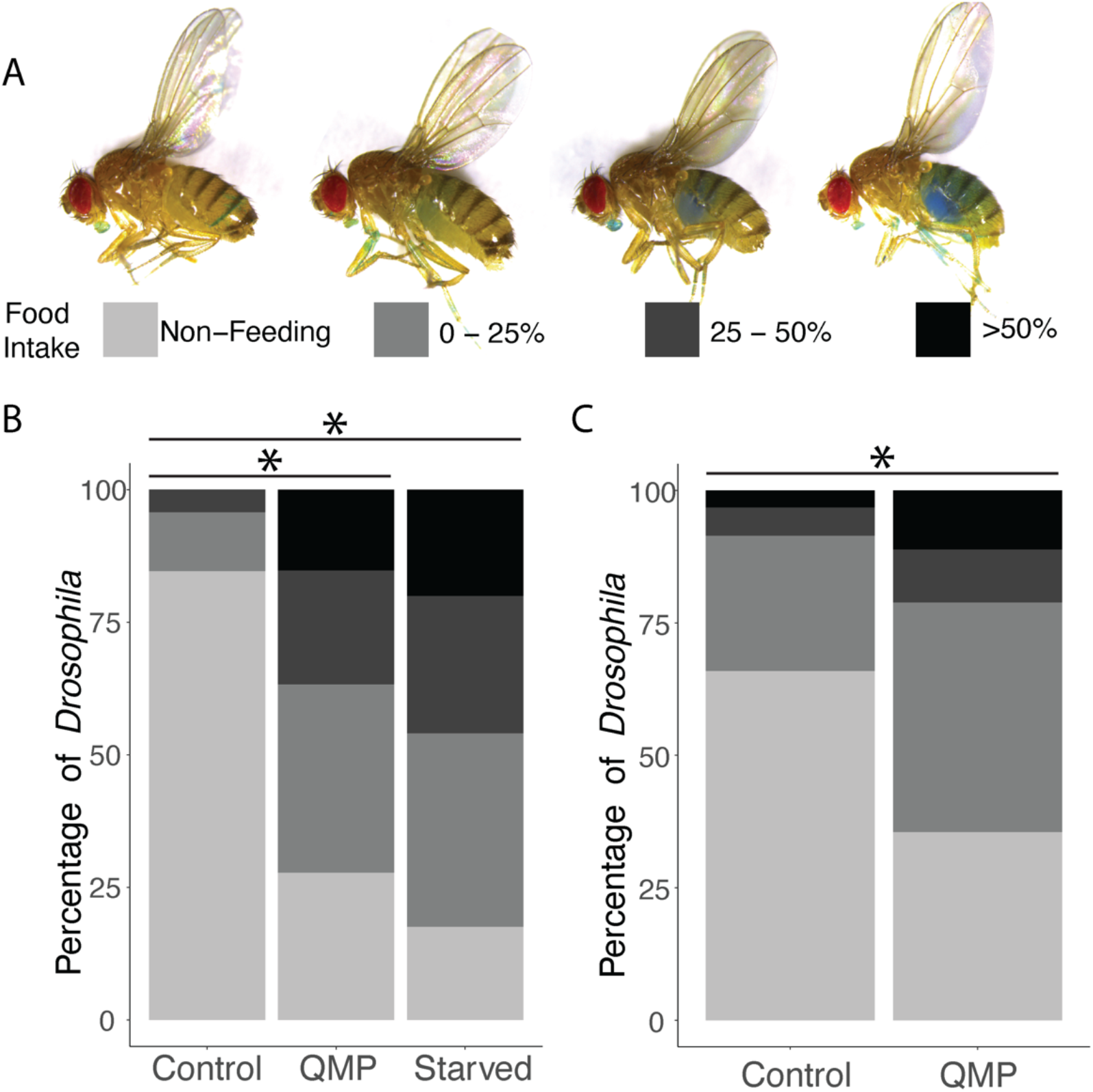
A) Representative images of feeding assays showing categorisation of food intake based on assays described in (Jiang et al., 2018). Food intake is categorised based on percentage of the abdomen which showed the presence of coloured food. B) Food consumption in *D. melanogaster* exposed to 26 Qe QMP or a solvent control while on a standard liquid diet for 48 hours or starved for 24 h prior to exposure to solvent control. **C)** Food consumption in *D. melanogaster* exposed to 26 Qe QMP or a solvent control while on a standard liquid diet for only 12 hours. A Fisher’s exact test was carried out, with significance being determined by a *p* value of < 0.05.

After 48 h of QMP exposure, an increase in food consumption was observed (Fig. 3B). Only 15% of the control population consumed food, whereas 74% of the QMP exposed population fed (*p* = 2.4 × 10^−17^), this is despite there being no restriction in access to food up to the start of the assay, as they had been fed standard solid media (see Methods, *Drosophila* stocks and maintenance). (Fig. 5). When this was further examined, the controls that did consume food mostly filled less than 25% of their abdomen, whereas half the QMP exposed filled more than 25% of their abdomen (*p* = 1.02 × 10^−13^) (Fig. 3B). This implies that QMP exposure not only increases the likelihood an individual will feed, but also increases the quantity of food that individual flies consume.

Given the difference in food consumption observed between the control and QMP treated flies (Fig. 3B), we hypothesised that QMP exposed individuals may be increasing intake in response to a perceived nutritional deficit as a response to QMP. We therefore tested if QMP treatment induced a feeding behaviour comparable to starvation. *D. melanogaster* were exposed to QMP or a solvent control while on the standard liquid diet for 48 h, before being starved for 24 h. The same food intake assay was carried out. Compared to the 15% of the fed control population (Fig 3B) which consumed food, 82% of the starved controls ate. When the fed individuals exposed to QMP were compared to the starved controls, there was no significant difference in food intake (*p* > 0.05) (Fig.3C). The quantities of food consumed by fed QMP and starved controls also were not different (*p* > 0.05) (Fig.3C). This indicates that QMP exposure is inducing a starvation like phenotype in exposed flies.

### The starvation feeding phenotype of QMP is established prior to ovarian repression

We have established that 48 hours of QMP exposure induced starvation-like feeding behaviour (Fig. 3B). Also, after 48 hours of QMP exposure, there is a clear reproductive repressive phenotype already established within the ovary in the 2a/b checkpoint, stage 9 checkpoint, and the number of mature oocytes (Figs. 2, B, D and 1B respectively). However, to determine if the differences in feeding (and thereby perception of nutritional status) are upstream of ovarian repression or if ovarian repression likely stimulated the differences in feeding response, we conducted the food consumption assay after only 12 h of QMP exposure. As shown in Figure 1B and 2B and D, there is no ovarian repression established in QMP treated flies by 12 hours of treatment. However, after only 12 h of QMP exposure, a significant difference in food intake was already established (Fig. 3C). The QMP exposed population was more likely to consume food than the controls (p = 9.09 × 10^−5^). Of the QMP exposed group, 64% of the population ate, compared to 34% of the controls. QMP therefore induces an increase in food intake prior to changes in ovarian phenotype, indicating that QMP is acting to alter nutrient sensing, or perception of nutrient status, and that this response is, at least partly, responsible for QMP mediated ovarian repression.

## Discussion

We have shown that repression of reproduction by QMP in *D. melanogaster* is likely acting through nutrient signalling, with honeybee QMP inducing sustained ovarian repression similar to that seen with starvation or nutrient deprivation (Fig. 3, (Burn et al., 2015; Pritchett and McCall, 2012; Pritchett et al., 2009)). Individuals exposed to QMP exhibit a significant increase in food consumption, which is indistinguishable from that caused by starving *D. melanogaster*. QMP appears to induce a perceived state of nutritional deficit, leading to increased food intake. This perceived nutritional deficit is established prior to any phenotype of repression being observed in the ovary-indicating that this is an upstream sensing or signalling process and that QMP may be acting on either nutrient sensing, hunger, or satiety signalling to induce this response (Lin et al., 2019). When an ovarian response is observed, this phenotype is also consistent with starvation. The 2a/b checkpoint in the germarium is activated, leading to a reduction in the number of oocytes which leave this region of the ovary-reducing reproductive potential. Alongside this, the stage 9 checkpoint in the ovary is activated, which induces degradation of stage 9 oocytes. Both of these are characteristic of starvation (Pritchett et al., 2009). Combined, these have a significant impact on the fecundity of the *D. melanogaster* exposed to QMP.

That QMP phenocopies or induces a starvation response in *D. melanogaster* implies that QMP interferes with nutrition signalling, or perception, in this species. That this effect is not limited to *D. melanogaster* (Camiletti et al., 2013; Lovegrove et al., 2019; Sannasi, 1969), but also occurs in a wide range of arthropods (Carlisle and Butler, 1956; Hrdy et al., 1960; Nayar, 1963; Princen et al., 2020; Sannasi and George, 1972) implies that whatever is being affected by QMP is conserved and present broadly in arthropods. It therefore seems likely that QMP is disrupting highly conserved nutrition perception or signalling pathways in these species. For example: QMP may disrupt include neuronal integration of gustatory perception, target-of-rapamycin or insulin signalling (Itskov and Ribeiro, 2013) or juvenile hormone metabolism; mechanisms which are known to link nutrition and reproduction in most insect species (Rodrigues and Flatt, 2016; Smykal and Raikhel, 2015).

Nutrition and nutrient sensing pathways have been implicated in establishing and maintaining reproductive skew in several social and eusocial arthropods (Kapheim, 2017). In particular, nutrient sensing pathways, specifically insulin signalling, are associated with social signals and reproductive state in ants (Chandra et al., 2018; Okada et al., 2017). In the eusocial paper wasp *Mischocyttarus mastigophorus*, there is evidence that the dominant most-reproductively active individuals are those which receive the highest levels of nutrition (Markiewicz and O’Donnell, 2001; O’Donnell et al., 2018), a trend which has been also shown in the wasp *Polistes dominulus* (Tibbetts, 2007) and *Polistes metricus* (Toth et al., 2009). Social spider colonies have their reproductive rates skewed in response to their nutritional environment, with individuals receiving a higher quality diet being the most reproductively active (Salomon et al., 2008). Genes which differ with caste, and therefore reproductive potential, in the bee *Megalopta genalis* have been shown to be associated with metabolism (Jones, Beryl et al., 2017). Nutrition is also associated with reproductive rate and reproductive diapause in solitary species (Mirth et al, 2019; Ojima et al., 2018), raising the possibility that QMP, and possibly other queen pheromones, have evolved to co-opt ancient mechanisms involved in environmentally responsive reproduction. Does then the nutrient sensing-reproductive repression mechanism induced by QMP in *Drosophila* reflect what is happening in honeybees?

Our findings in *Drosophila* are superficially consistent with what is known in the honeybee ovarian response to QMP. In honeybees, QMP acts within the germarium to regulate oogenesis (Duncan et al., 2016) as it does in *D. melanogaster* through activation of the 2a/b checkpoint. QMP in honeybees is also suggested to act to induce apoptosis at later stages of oocyte maturation (Ronai et al., 2015), and we have shown the degeneration of stage 9 oocytes in *D. melanogaster* in response to QMP (Fig. 2). Importantly, however, a key effect of QMP on *D. melanogaster* is not paralleled in honeybees; the increase in food consumption QMP causes in *D. melanogaster*. QMP exposed, 4 day old worker bees are more able to resist starvation than those not exposed to QMP, a phenotype linked to higher lipid stores in their fat body (Fischer and Grozinger, 2008), implying a link between starvation phenotypes and QMP in honeybees. Honeybee experiments comparing worker bees across the first 10 days after emergence with QMP and those without QMP, however, show no differences in food consumption between treated and controls (Duncan et al., 2016, 2020). This implies that if, as we propose here, an ancient ‘arthropod-wide’ mechanism linking nutrition with reproduction has been co-opted in the evolution of QMP, that this link between nutrition and reproduction has been decoupled in the honeybee lineage. Consistent with this, there have been significant changes in the regulatory interactions linking nutrition, neuroendocrine signalling and reproduction in honeybees and other eusocial insects (Rodrigues and Flatt, 2016; Kapheim, 2017).

Our data links QMP with nutrition signalling in flies, and perhaps, by extension, the broad range of species where QMP has a similar effect (Carlisle and Butler, 1956; Hrdy et al., 1960; Nayar, 1963; Sannasi, 1969). That nutrition signalling is implicated in eusociality across a broad range of arthropods, and multiple independent evolutions of eusociality, implies a deep role for these pathways in the evolution of eusociality. We propose that these data point to ‘queen pheromones’ evolving to implement the reproductive division of labour through the manipulation of nutrient sensing pathways in non-reproductive individuals (Okada et al., 2017; Toth, 2017).

QMP is a highly derived pheromone, consisting of five major semiochemicals (Slessor et al., 1988) that are chemically distinct from the less derived and more broadly used queen pheromones from other eusocial insects (Van Oystaeyen et al., 2014)(predominantly linear alkanes). These less-derived queen pheromones do not induce ovary repression in *D. melanogaster* (Lovegrove et al., 2019), suggesting that QMP may be unique in its ability to control reproduction in non-target species. Given the chemical complexity of QMP, that it has evolved over the last 55 million years (Peters et al., 2017), the broad range of the response to QMP outside eusocial insects (Carlisle and Butler, 1956; Hrdy et al., 1960; Nayar, 1963; Sannasi, 1969) (Figure 4), and the data we present here linking QMP with manipulation of nutrient sensing in *Drosophila* we propose that QMP has evolved, as a result of an evolutionary ‘arms race’ over worker reproduction, to target deeply conserved essential and pleiotropic pathways such as Notch signalling (Duncan et al., 2016) and neuroendocrine signalling. It has not escaped our notice that this interpretation would suggest manipulative (Strauss et al., 2008), rather than honest (Keller and Nonacs, 1993), signalling of reproductive state in the case of QMP.

Our discovery that QMP triggers a starvation response in *D. melanogaster* allows us to link queen pheromones to nutrition sensing. This link, between a recently evolved pheromone and presumably ancient, highly conserved nutrient sensing pathway may underlie the repeated evolution of eusociality, providing a route to a most successful life-history strategy; eusociality.

## Materials and Methods

### D. melanogaster stocks and maintenance

*D. melanogaster* used in this study were maintained at 25 °C on a 12h:12h light/dark cycle. All flies were Oregon-R modENCODE line (Stock #25211) from the Bloomington *Drosophila* stock centre. They were reared on a solid yeast/sugar medium of; 3 L dH_2_O, 200 g organic cornmeal, 50 g brewer’s yeast, 140 g sugar, 20 ml propionic acid and 15 ml 10% methyl *p*-hydroxybenzoate in absolute ethanol. Virgin females were anaesthetised with CO_2_ collected within one hour of emergence. Virgin females had meconium present within an enlarged abdomen, and an overall pale colouration and were stored separately from the parent population, at room temperature for 24 h prior to use.

### QMP dilution

QMP is measured in Queen equivalents (Qe), with one Qe being the amount a mated queen will produce in a 24 h period (Pankiw et al., 1996). QMP consists of five major semiochemicals (Slessor et al., 1988). One Qe for a European mated queen *Apis mellifera* contains; 200 μg 9-keto-(E)-2-decanoic acid (ODA), 80 μg 9-hydroxy-(E)-2-decanoic acid (9-HDA) and 20 μg methyl-hydroxybenzoate (HOB) and 2 μg 4-hyroxy-3-methoxyphenylethanol (HVA) (Pankiw et al., 1996). QMP (Intko Supply Ltd, Canada) was dissolved in absolute ethanol to a concentration of 26 Qe/20 μl, and stored at -20 °C until use.

### QMP exposure

QMP exposure was carried out in modified 50 ml centrifuge tubes as previously described (Lovegrove et al., 2019). *D. melanogaster* were fed 500 μl of liquid diet per day (made in batches of 5 ml on the day of use; 4.75 ml dH_2_O, 5% absolute ethanol, 0.15 g sugar and 0.1 g brewer’s yeast (Camiletti et al., 2013)). For starvation conditions, the diet was 500 μl of 5% ethanol solution in dH_2_O (no sugar or yeast). On top of either of these diets, 20 μl of 26 Qe QMP or 20 μl of ethanol solvent control was added. The 24 h old virgin females (n = 10 per vial, each treatment having at least 5 replicates) were anesthetised with CO_2_ and added to the tube, and the end blocked with a cotton ball. The tube was left on its side until all individuals had recovered from CO_2_ narcosis. *D. melanogaster* were incubated at 25 °C for 12 -48 hours (trial dependent).

### Ovary collection and fixation

*D. melanogaster* were anesthetised with CO_2,_ and ovaries dissected into ice-cold PBS (phosphate buffered saline). Ovaries which were damaged, or lost mature oocytes, were discarded. Ovaries were fixed using 4% formaldehyde in PBS at room temperature for 10 min. Ovaries were washed 4x with 1 ml of PTx (0.1% Triton-X100 in PBS). During the last wash, 1 μl of DAPI (4′,6-diamidino-2-phenylindole) was added, and incubated in the dark 15 min. Ovaries were washed with PTx twice more before being stored at 4 °C in the dark in 70% ultrapure glycerol for >24 h prior to mounting (as described in (Lovegrove et al., 2019)). Ovaries were bridge mounted in 70% ultrapure glycerol before the number of stage 14 mature (vitellogenic) oocytes were counted manually under a Leica L2 dissection microscope. This was used as an indicator of fecundity (King, 1970). DAPI staining was visualised under an Olympus BX61 Fluoview FV100 confocal microscope with FV10-ASW 3.0 imaging.

### Food intake assay

To visualise the food intake of *D. melanogaster*, liquid diets were created. These were 5% brewer’s yeast and 5% sugar (dissolved) in dH_2_O. To these food colouring was added to a final concentration of 5%. These were aliquoted and stored at -20 °C until use. Virgin females were exposed to 26 Qe QMP or a solvent control for 12 or 48 h. Also included was a group which had been exposed to the starvation liquid diet for 48 hours, along with the solvent control. They were then transferred to a test plate for the feeding trial. This was a petri dish containing 3% agar, on top of which 3 x 100 μl drops of yeast solution were alternated around the edge of the plate with 3 x 100 μl of sugar solution. The *D. melanogaster* were anesthetised with CO_2_ before being transferred to the plate, and incubated at room temperature until they had recovered. The plates were incubated in the dark for 2 hours at 25 °C. After 2 h they were frozen at – 20 °C to prevent further feeding. This was carried out on 10 vials of 10 individuals for QMP treated and controls.

Quantification of food intake was determined by visually inspecting each *D. melanogaster* after freezing and looking for evidence of coloured food within their abdomen. They were classified based on the scale described in (Jiang et al., 2018). Individuals which had not consumed food were scored a 0. Those which had consumed enough to only lightly colour their abdomen or fill less than 25% were classified as 1. Those which had darker abdomens and had filled 25 – 50% of their abdomens were scored as 2, and those which filled over 50% of their abdomen were classified as 3.

### Determination of ovarian checkpoint activation

During oocyte development and maturation in *D. melanogaster*, there are checkpoints along which reproduction may be repressed in adverse environmental or nutritional conditions (Pritchett et al., 2009). The first of these is located in the germarium at stage 2a/b, the activation of which reduces the output of oocytes for maturation, effectively slowing the overall rate of reproduction (Drummond-Barbosa and Spradling, 2001). The second of these is located at stage 9 in mid-oogenesis, resulting in the degradation and death of stage 9 oocytes (McCall, 2004) (Fig. 2A).

*D. melanogaster* oocytes were staged by visualising DAPI staining (Jia et al., 2016; King, 1970)(Fig. 2A). The number of germaria, healthy stage 9 oocytes, degenerating stage 9 oocytes and stage 10 oocytes were counted. Stage 9 oocytes were characterised based on the beginning of migration of anterior follicle cells and stage 10 was classified as occurring after the migration of the anterior follicle cells (Jia et al., 2016; King, 1970). Stage 9 degradation was observed as a loss of structural integrity, coupled with bright, fragmented nuclei (Fig. 2A).

Evidence of early ovarian checkpoint activation in response to QMP was determined by calculating the ratio of the number of oocytes which reach or successfully pass the second checkpoint (stage 9 (S9), stage 9 degenerating (S9d), stage 10(S10) per germaria (e.g., containing the germline stem cell niche). This can be thought of as the theoretical versus actual reproductive capacity of an individual.

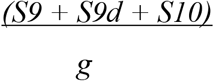

If every germarium is producing presumptive oocytes which will pass through the 2a/b ovarian checkpoint, and progress to the stage 9 checkpoint we would expect a ratio of approximately 1:1, shown numerically as 1.0. However, if the stage 2a/b checkpoint is activated then the number of oocytes passing through to the stage 9 checkpoint will be reduced, lowering reproductive output below that of the anticipated theoretical value. This is reflected in a ratio significantly lower than 1.0. For example, if the 2a/b checkpoint activation reduces the number of oocytes reaching that stage 9 checkpoint by 50%, then the ratio will be 0.5.

### Calculation of stage 9 oocyte checkpoint activation

The proportions of stage 9, stage 9 degenerating and stage 10 oocytes were calculated from count data. Any activity of the stage 9 checkpoint should lead to an increase in the proportion of oocytes at stage 9 which show degradation. This is likely to also be associated with a reduction of oocytes which pass through this checkpoint at stage 9, and successfully reach stage 10.

### Statistical analysis

Data was analysed using R Studio version 1.2.5033 running R version 3.6.2. Assessment of whether the data fit a normal distribution was carried out using a Shapiro-Wilk test, all data showed a non-normal distribution and so data was analysed using Generalised Linear Mixed Models (GLMMs) using lme4 (Bates et al., 2015). In all cases treatment was treated as a fixed effect and the slide number as a random factor. The maximal model was simplified using Analysis of Deviance (AOD) to assess the effect of removing terms. Where an effect of treatment was found, pairwise comparisons between treatments were carried out using emmeans using a Tukey post-hoc test, to correct for multiple testing. Ovariole numbers (Fig. 1A) were analysed using a Poisson error structure with a log link. The number of mature oocytes (Fig. 1B) were analysed using a negative binomial error structure after model fitted with the Poisson error structure was shown to have higher than expected residual variance (over-dispersed). The ratio of oocytes reaching the second ovarian checkpoint was analysed with a Gaussian error structure (Fig. 2A). Because we expect the numbers of ovarioles, stage 9 oocytes, degenerating oocytes and stage 10 oocytes to be dependent variables, we analysed the data in Fig. 2B by performing a principle components analysis (PCA), which indicated that the majority of variation was observed in the first principle component. This component was extracted and a GLMM with a Gaussian error structure was used to determine the effect of treatments at different time points. Analysis to determine whether there were differences in food consumption (Fig. 3) were assessed using a Fisher’s exact test.

## Author contributions

MRL: Assisted with experimental design, carried out *D. melanogaster* experiments and statistical analysis (Fig. 3) drafted and edited manuscript.

EJD: Assisted with experimental design, carried out statistical analysis (Fig. 1, 2), assisted with preparation of figures and edited manuscript.

PKD: Assisted with experimental design, supervised *Drosophila* experiments, assisted with preparation of figures and drafted and edited manuscript.

## Acknowledgements

MRL was funded by a University of Otago Doctoral scholarship. EJD was supported by a Marie Skłodowska-Curie Individual Fellowship (H2020-MSCA-IF-2016 752656). The authors thank Dr Jens van Eeckhoven and Dr James Rouse for statistical advice.

